# Engineering of extracellular vesicles for small molecule-regulated cargo loading and cytoplasmic delivery of bioactive proteins

**DOI:** 10.1101/2021.11.04.466099

**Authors:** Masaharu Somiya, Shun’ichi Kuroda

## Abstract

Cytoplasmic delivery of functional proteins into target cells remains challenging for many biological agents to exert their therapeutic effects. Extracellular vesicles (EVs) are expected to be a promising platform for protein delivery; however, efficient loading of proteins of interest (POIs) into EVs remains elusive. In this study, we utilized small compound-induced heterodimerization between FK506 binding protein (FKBP) and FKBP12-rapamycin-binding (FRB) domain, to sort bioactive proteins into EVs using the FRB-FKBP system. When CD81, a typical EV marker protein, and POI were fused with FKBP and FRB, respectively, rapamycin induced the binding of these proteins through FKBP-FRB interaction and recruited the POIs into EVs. The released EVs, displaying virus-derived membrane fusion protein, delivered the POI cargo into recipient cells and their functionality in the recipient cells was confirmed. Furthermore, we demonstrated that CD81 could be replaced with other EV-enriched proteins, such as CD63 or HIV Gag. Thus, the FRB-FKBP system enables the delivery of functional proteins and paves the way for EV-based protein delivery platforms.

## Introduction

Current biopharmaceutics, such as nucleic acid therapeutics, RNA-based vaccines and therapeutics, and genome editing are heavily dependent on the development of precision delivery technologies^1–3^. As the active pharmaceutical ingredients (APIs) of biopharmaceutics are generally large biomolecules, such as DNA, RNA, and proteins, they cannot penetrate the cell membrane and are thus incapable of exerting therapeutic efficacy without a delivery system. The primary purpose of a delivery system for biotherapeutics is to protect the APIs from degradation following uptake by target cells, facilitate penetration of the cell membrane, and aid in reaching the cytoplasm.

One of the most successful delivery systems for biomolecules is the lipid nanoparticle (LNP) platform, which is designed to deliver nucleic acid therapeutics, including small interfering RNA and mRNA^4^. After injection into the body, LNPs are internalized into cells by endocytosis, and because of the low pH-dependent protonation properties of an ionizable lipid, LNPs can escape from the endosomal compartment and thus deliver their nucleic acid cargo into the cytoplasm. Despite their superior delivery competence, current LNPs can only target the liver cells after systemic injection or the local immune cells after intramuscular injection. For broader targeting of cells other than liver and local immune cells, the development of an alternative delivery system for functional biomolecules is in high demand.

Unlike the LNPs for nucleic acid therapeutics, no feasible platform has been established for intracellular protein delivery despite decades of basic research. One of the main reasons that has hampered the translation of protein therapeutics into the clinic is the inefficiency of the endosomal escape of protein delivery systems. Currently available delivery systems for proteins include lipid-based nanoparticles, nano-micelles, and inorganic nanomaterials^5^. The endosomal escape efficacy of these delivery systems is still far from sufficient for clinical applications^6^. An ideal platform for intracellular protein delivery should possess the ability to encapsulate the protein cargo, be taken up by target cells, and efficiently escape the endosomal compartment, to release the protein cargo into the cytoplasm.

Extracellular vesicles (EVs) are considered a novel modality for the delivery of biomolecules^7^. They are produced by cells and are thus considered safe delivery systems capable of delivering large biomolecules, such as proteins and nucleic acids, because they can encapsulate any materials that cells produce. Furthermore, the engineering of EVs can exploit novel functionalities to achieve better pharmacokinetics and intracellular delivery by expressing functional proteins in EV-producing cells^8^.

Despite the huge potential of EVs as drug delivery systems, their application still has some limitations. One of the most intractable issues is that there is no efficient method to actively load therapeutic molecules into EVs^9^. Furthermore, the low efficacy of non-modified EVs for endosomal escape needs to be improved using an engineering approach. Our recent work demonstrated that non-modified EVs failed to deliver their protein cargo into recipient cells, whereas conjugation with virus-derived membrane fusion protein significantly improved the delivery efficiency^10,11^.

In this study, we developed a controlled loading method for specific proteins in EVs, which is regulated by a compound. In the presence of rapamycin, a compound that induces heterodimerization of FK506 binding protein (FKBP) and FKBP12-rapamycin-binding (FRB) domains, the protein of interest (POI) was efficiently encapsulated into EVs and secreted into the extracellular space of donor cells. Engineered EVs harboring virus-derived fusogenic proteins can deliver encapsulated POI into the cytoplasm of recipient cells.

## Materials and Methods

### Materials

The compounds, NanoLuc substrate, antibodies, and other materials used in this study are listed in Supplementary Table 1. The plasmids used in this study were constructed using a conventional PCR-based method^12^. Supplementary Table 2 lists the plasmids used in the present study.

### Cell culture, transfection, and EV preparation

Human embryonic kidney HEK293T cells (RIKEN Cell Bank) were cultured in Dulbecco’s modified Eagle medium (DMEM, high glucose formulation, Nacalai Tesque) with 10% fetal bovine serum (FBS) and 10 μg/mL penicillin-streptomycin at 37°C with 5% CO_2_ in a humidified atmosphere.

HEK293T cells were transfected using 25-kDa branched polyethyleneimine (PEI, Sigma) as previously described^10^. Briefly, plasmid DNA and PEI were mixed in Opti-MEM (Thermo Fisher) and applied to the cells that were seeded one day prior to transfection (e.g., 10^6^ cells/dish in 60 mm-dish). The weight ratio of plasmid DNA to PEI was 1:4. The plasmid formulation for the transfection of EV donor cells consisted of the following: FKBP-fusion protein:FRB-POI:VSV-G = 2:2:1 (weight). After transfection, cells were cultured for 24–96 h in the presence or absence of rapamycin (1–100 nM, final concentration), and the supernatant and cells were harvested and subjected to the following experiments.

EVs in the culture supernatant were concentrated by PEG precipitation, as described previously^10^. Typically, 5 mL of supernatant from 3–4 days of cell culture was concentrated to 100 μL (50-fold concentration) in phosphate buffered saline (PBS) and used immediately for the reporter gene assay.

### Characterization of EVs

Protein expression in the donor cells and EVs was analyzed by western blotting as described previously^10^. To evaluate the loading efficiency of HiBiT-tagged POIs into EVs, the HiBiT content in the concentrated EV was estimated from the measurement of NanoLuc activity by mixing with NanoLuc substrate and LgBiT according to the manufacturer’s instructions. The luminescence signal was quantified using a plate reader (Synergy 2, BioTek).

The loading efficiency of POIs into EVs was evaluated by co-immunoprecipitation^10^. The culture supernatant was clarified by low-speed centrifugation (1,500 ×g, 5 min) and mixed with Protein G magnetic beads coated with mouse monoclonal anti-human CD81 or mouse IgG antibodies for over 20 min. After washing with PBS at least four times, the magnetic beads were mixed with LgBiT and NanoLuc substrate (Nano Glo^®^ HiBiT Lytic Detection System), and luminescence signals were measured.

### Luminescence reporter gene assay for functional delivery of POIs

The delivery efficiency of functional FRB-tTA protein was evaluated using a reporter gene assay as described previously^11^ with a slight modification. Recipient HEK293T cells (approximately 2×104 cells/well in a 96-well plate) were transfected with plasmids encoding NanoLuc with PEST sequence^13^ under the TRE promoter the day before treatment with EVs. Unless otherwise indicated, the recipient cells were treated with 10 μL of concentrated EVs, which was equivalent to 0.5 mL of parental cell supernatant (this is based on the estimation that supernatant was approximately 50-fold enriched by PEG precipitation) for 24 h. After treatment with EVs, the recipient cells were lysed and mixed with NanoLuc substrate according to the manufacturer’s instructions, and luminescence signals were measured. Induction of the reporter gene expression was expressed as fold-change of EV-treated cells compared to non-treated cells, that is, the observed luminescence signal was normalized to that of non-treated cells.

### Fluorescence reporter gene assay

The functional delivery of FRB-Cre-HiBiT was evaluated using a reporter gene assay described previously^11^. Briefly, HEK293T cells (approximately 2×10^4^ cells/well in a 96-well plate) were transfected with a reporter plasmid (encoding LoxP-flanked mKate and EGFP under the CMV promoter), treated with EVs for 24 h, and then observed under a fluorescence IX70 microscope (Olympus).

### Statistical analysis

The data were statistically analyzed by Student’s *t*-test or one-way ANOVA followed by either *post hoc* Tukey’s HSD or Dunnett’s test using the Real Statistics Resource Pack software created by Charles Zaiontz.

## Results

### Controlled loading of bioactive proteins into EVs

Controlling the loading of specific proteins into EVs remains challenging. We first tried to establish controlled loading of POI into EVs by well-characterized small compound-dependent protein dimerization, based on a similar approach reported in 2019^14^. FKBP and the FRB domain of mTOR form a heterodimer in the presence of rapamycin or its analog^15,16^. We expected that this compound-mediated heterodimerization could induce the recruitment of specific proteins to EVs. In this study, a typical EV marker protein, CD81, was fused with FKBP at the C-terminal cytoplasmic tail, and the POI was fused with FRB at the N-terminus (Fig. 1, upper panel). These proteins can induce the interaction of CD81 and POI via FKBP-FRB heterodimerization in the presence of rapamycin (Fig. 1, lower). In this study, we utilized two POIs, FRB-tTA and C-terminally HiBiT-tagged FRB-Cre protein, where the HiBiT tag enables the precise quantification of tagged proteins by luminescence assay^10,17^.

**Fig. 1.**
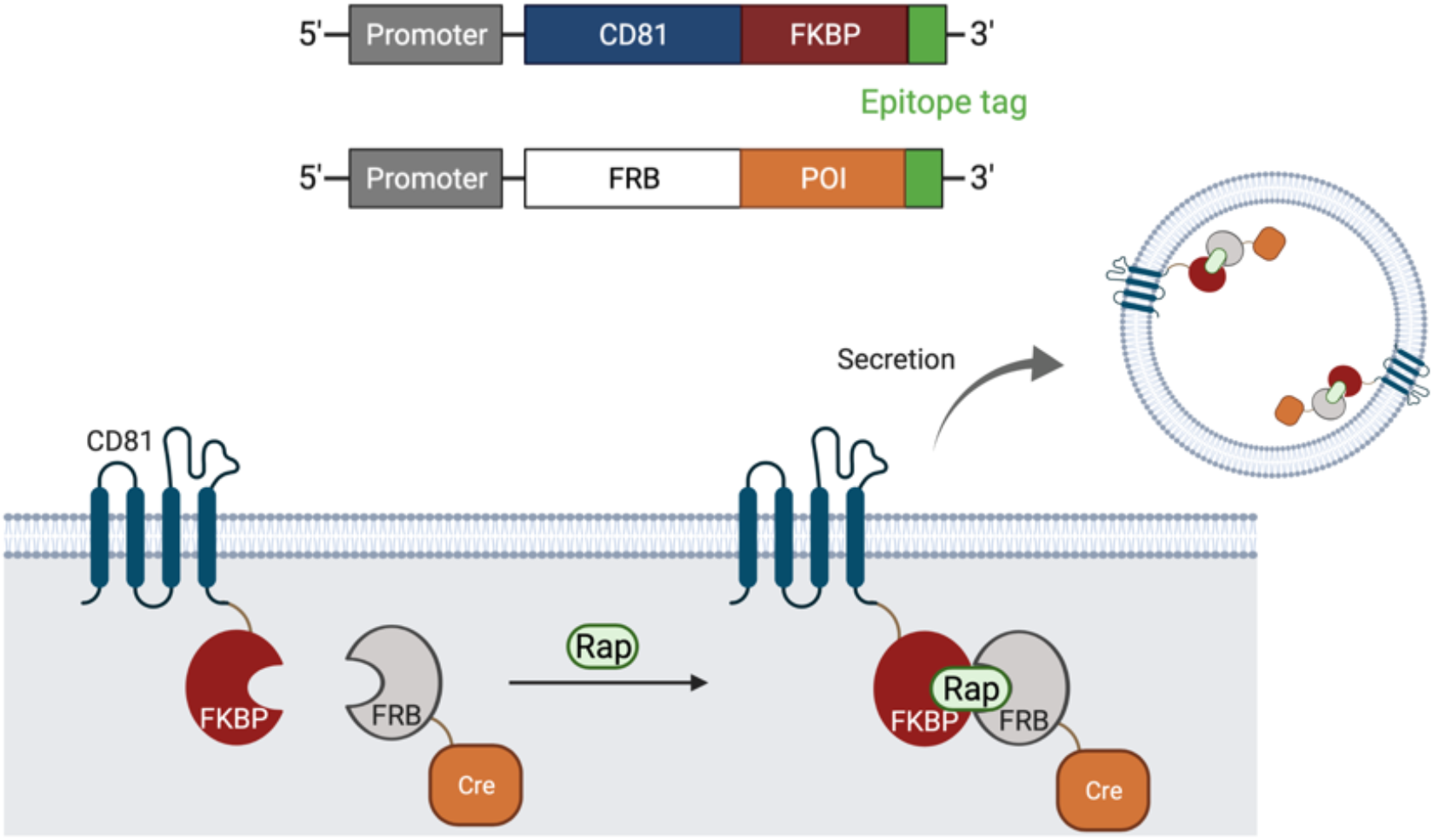
Schematic representation of rapamycin-induced recruitment of POIs into EV following FKBP-FRB heterodimerization.

We transfected donor HEK293T cells with plasmids encoding CD81-FKBP, FRB-POI, and VSV-G, a viral fusogenic protein that is known to enhance the delivery efficiency of EVs^10,18^. Protein expression was analyzed by western blotting (Fig. 2A), and all proteins were found to be expressed well in the HEK293T cells. After culturing in the presence or absence of rapamycin, we collected the supernatant of transfected cells and measured the amount of POI (FRB-tTA or FRB-Cre-HiBiT) in the concentrated EVs. Western blotting analysis revealed that all expressed proteins (CD81-FKBP, FRB-tTA, and VSV-G) were found in the donor cell lysate (Fig. 2B). CD81-FKBP and VSV-G were also found in the concentrated EVs. As expected, EV-negative marker calnexin was not seen in the EV samples. FRB-tTA was found in the concentrated EVs when the donor cells were cultured in the presence of rapamycin, suggesting that FRB-tTA was loaded into the EVs and secreted. The amount of FRB-Cre-HiBiT in concentrated EVs was significantly increased as the concentration of rapamycin increased, suggesting that FRB-Cre-HiBiT binds to CD81-FKBP via rapamycin and is released from cells along with EVs (Fig. 2C).

**Fig. 2.**
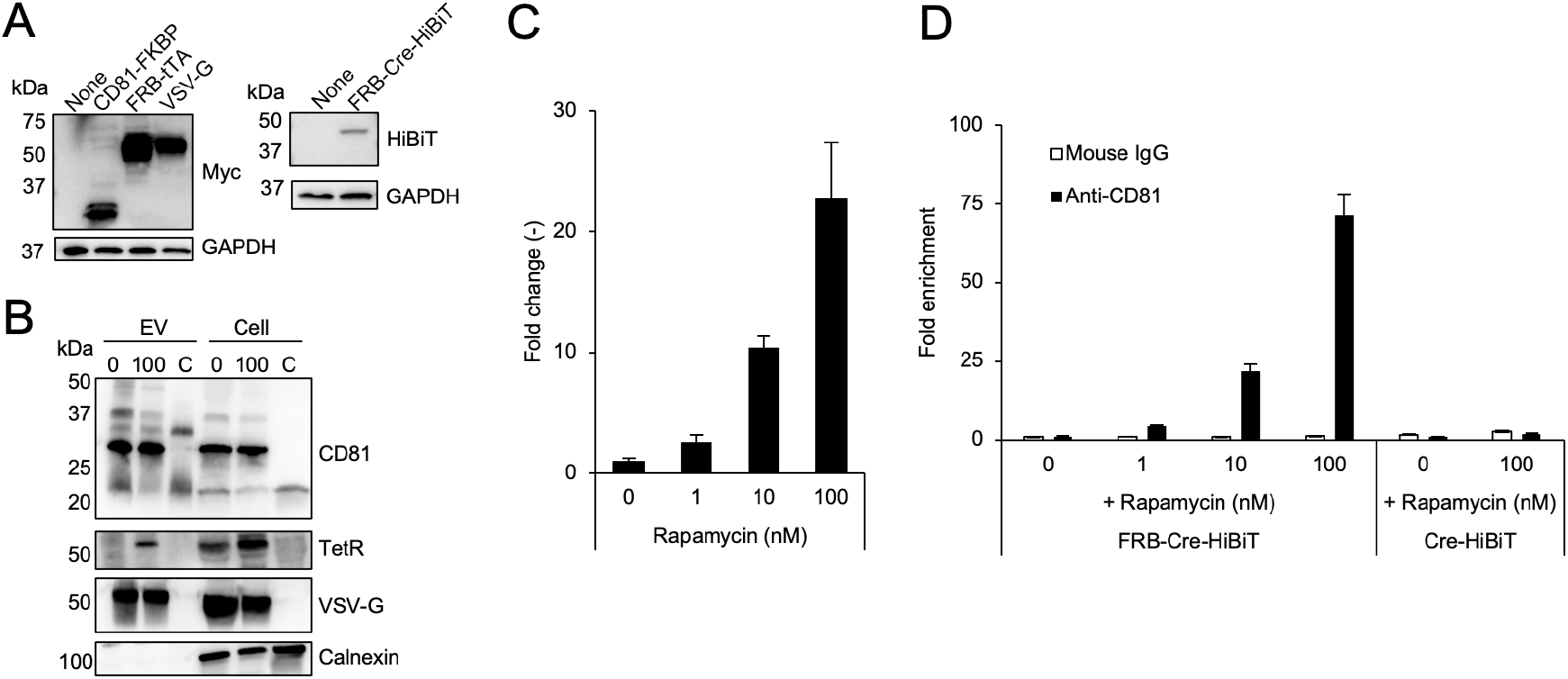
Rapamycin-induced recruitment of FRB-POIs into EVs expressing CD81-FKBP. (A) Protein expression in the donor HEK293T cells. Overexpression of proteins in cell lysates was detected by western blotting. The C-terminal Myc tag of CD81-FKBP, FRB-tTA, VSV-G were detected using an anti-Myc antibody. FRB-Cre-HiBiT was detected using an anti-HiBiT antibody. GAPDH served as a loading control. (B) Western blotting of concentrated EVs and donor cell lysate. Concentrated EVs or donor cell lysate (0, 100, and C represents rapamycin = 0 nM, 100 nM, and non-transfected control, respectively) were analyzed using anti-CD81, anti-TetR, anti-VSV-G, and anti-calnexin antibodies for the detection of CD81-FKBP, FRB-tTA, VSV-G, and calnexin, respectively. (C) Quantification of FRB-Cre-HiBiT in the concentrated EVs. The amount of FRB-Cre-HiBiT was calculated by measuring the HiBiT tag as a proxy. N = 3, means ± SD. (D) Co-immunoprecipitation of FRB-Cre-HiBiT using an anti-CD81 antibody. Culture supernatants from donor HEK293T cells expressing CD81-FKBP, FRB-Cre-HiBiT, and VSV-G were co-immunoprecipitated with an anti-CD81 antibody or total mouse IgG antibody as a control. The amount of FRB-Cre-HiBiT or Cre-HiBiT in the immunoprecipitate was measured by luminescence signal following addition of LgBiT and NanoLuc substrate. Fold-enrichment stands for the relative luminescence intensity normalized to that of the control sample (rapamycin = 0 nM). N = 3, means ± SD.

To demonstrate the encapsulation of FRB-Cre-HiBiT into the luminal side of EVs containing CD81-FKBP, culture supernatants from donor cells were immunoprecipitated with anti-CD81 antibody and the amount of FRB-Cre-HiBiT that immunoprecipitated together with the CD81-FKBP-containing EVs was measured. Compared to the control antibody (mouse IgG), anti-CD81 antibody efficiently co-immunoprecipitated FRB-Cre-HiBiT, and the amount of FRB-Cre-HiBiT increased depending on the concentration of rapamycin. Up to 70-fold enrichment of FRB-Cre-HiBiT was achieved in EVs by 100 nM rapamycin compared to the control. In contrast, without rapamycin, we observed an undetectable level of FRB-Cre-HiBiT, suggesting that passive loading of overexpressed FRB-Cre-HiBiT into EVs was almost negligible. Furthermore, without FRB fusion (Cre-HiBiT), no enrichment was observed following rapamycin treatment, indicating that FRB fusion is critical for loading the POIs into EVs (Fig. 2D).

### Functional protein delivery into recipient cells

Next, we evaluated the EV-mediated cytoplasmic delivery of POIs into recipient HEK293T cells. In this experiment, we used the previously described assay^11^ to measure the functional delivery of proteins, in this case, tetracycline transactivator (tTA). Following delivery of FRB-tTA into the cytoplasm of recipient cells, the reporter gene under the control of a TRE promoter is expressed, and the expression level of the reporter gene reflects the delivery efficiency.

Donor HEK293T cells were transfected with plasmids encoding CD81-FKBP, FRB-tTA, and VSV-G and cultured in the presence or absence of 10 nM rapamycin. After precipitation of the supernatant with PEG, the concentrated EVs from the donor cells were applied to the recipient HEK293T cells. We observed a significant increase in reporter gene expression in the recipient cells treated with EVs from donor cells cultured with rapamycin, whereas EVs from the no rapamycin condition barely induced any reporter gene expression (Fig. 3A). This result indicated that FRB-tTA was incorporated into the EVs in the presence of rapamycin and functional protein was delivered to the recipient cells. This result was consistent with the previous finding that passive loading of FRB-POI into EVs is not a significant portion of the cargo loading (Fig. 2D) and barely contributes to cargo delivery.

**Fig. 3.**
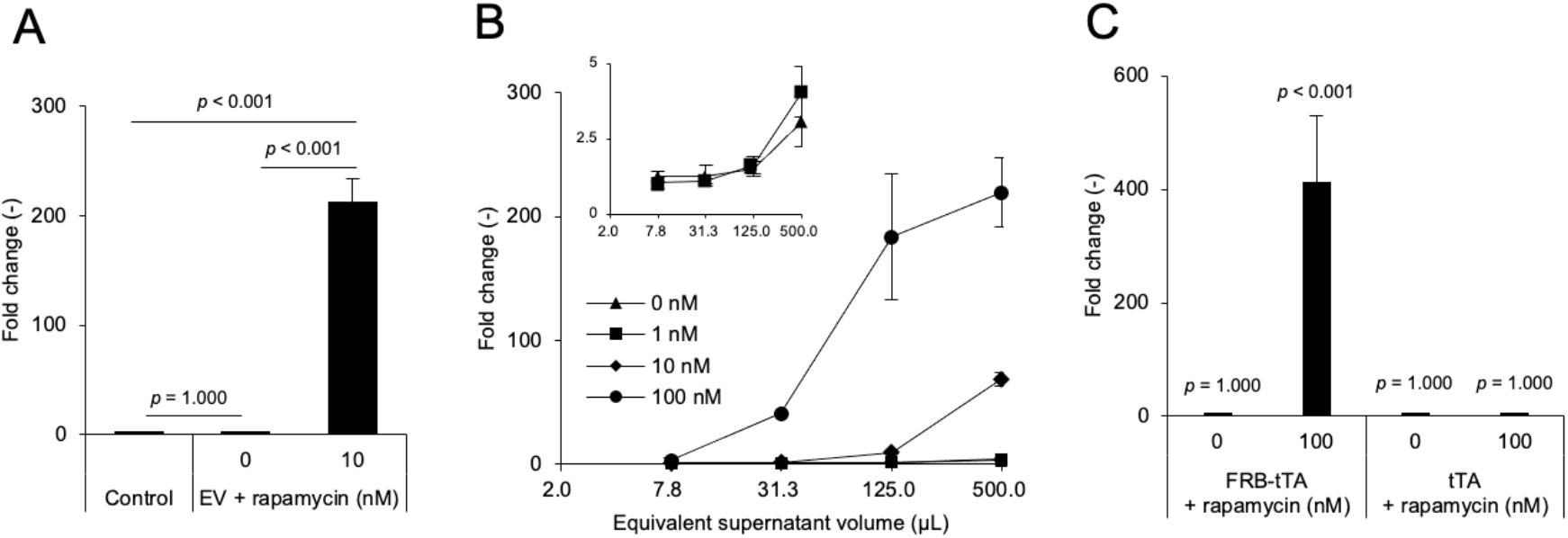
Reporter gene assay. Recipient HEK293T cells were treated with EVs containing CD81-FKBP, FRB-tTA, and VSV-G, and cultured for 24 h. Reporter gene expression was evaluated by measuring luminescence signal derived from the NanoLuc reporter. (A) Recipient cells were treated with EVs from donor cells cultured with 0 or 10 nM of rapamycin. N = 3, means ± SD. Data were analyzed by one-way ANOVA followed by Tukey’s HSD test. (B) Various doses of EVs from donor cells cultured with 0, 1, 10, and 100 nM of rapamycin were applied to the recipient cells. EV dose is represented as volume equivalent to the parental supernatant. The inset focuses on rapamycin concentrations of 0 and 1 nM. N = 3, means ± SD. (C) EVs from donor cells expressing either FRB-tTA or tTA were incubated with the recipient cells. N = 3, means ± SD. Data were analyzed by one-way ANOVA followed by Dunnett’s *post hoc* test against the non-treatment control.

Fig. 3B shows the clear dose-dependency of reporter gene expression by the EVs. As the rapamycin concentration increased during the culture of the donor cells, the resulting EVs induced stronger reporter gene expression. This may reflect the increased loading of FRB-POI into EVs in a rapamycin dose-dependent manner (Fig. 2B to 2D). Although the induction of reporter gene expression in the recipient cells by EVs from donor cells cultured under no rapamycin condition was not statistically significant, a slight induction (up to 5-fold) was observed at the highest EV dose (Fig. 3B, inset). This is likely due to the passive loading of a trace amount of FRB-tTA into EVs and their delivery into the cytoplasm of recipient cells. Furthermore, to confirm the FRB-mediated recruitment of tTA into the EVs, we transfected the donor cells with plasmids encoding CD81-FKBP, VSV-G, either FRB-tTA or tTA, and cultured the cells with or without rapamycin. The recipient cells were cultured with EVs, and reporter gene expression was measured (Fig. 3C). This experiment demonstrated that tTA without FRB fusion was not delivered to the recipient cells regardless of the rapamycin treatment of parental donor cells, suggesting that tTA without FRB domain was not actively recruited into the EVs, and that the fusion of the FRB domain was critical for the recruitment of POI into the EVs.

### Cargo protein delivery mechanism

As demonstrated previously, endosomal escape is the major rate-limiting step in EV-mediated intracellular drug delivery^10,11^. These studies described the enhancement of membrane fusion and endosomal escape using VSV-G. We confirmed that functional protein delivery using our system was driven by the fusogenic activity of VSV-G (Fig. 4A). When the donor cells were transfected with plasmids encoding a fusion-deficit mutant, VSV-G (P127D^19^) or a control protein (firefly luciferase, Fluc), no reporter gene expression was induced in the EV-treated recipient cells.

**Fig. 4.**
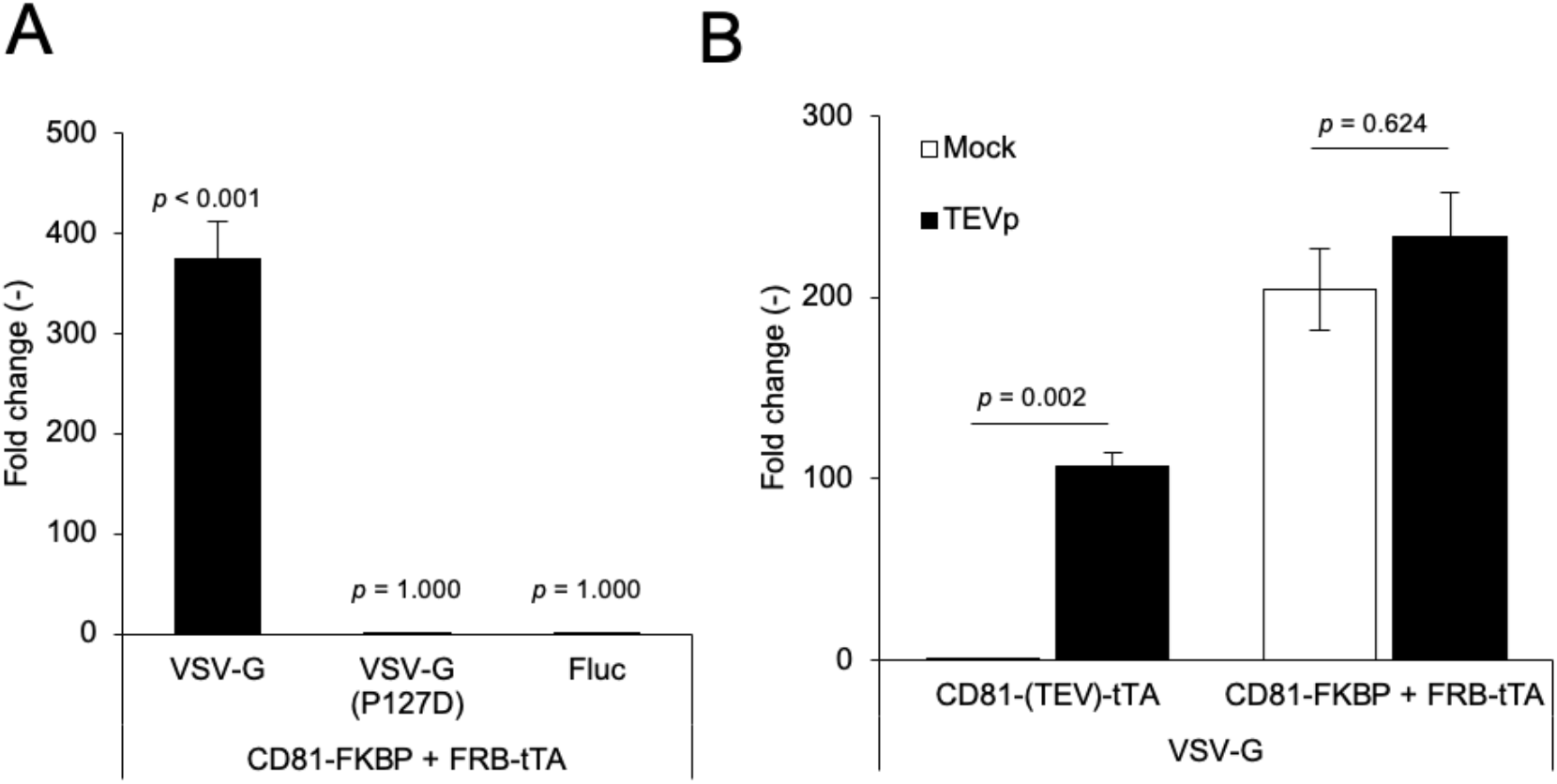
Mechanism of delivery of EVs containing CD81-FKBP, FRB-tTA, and VSV-G. (A) Contribution of the fusogenic activity of VSV-G in delivery of functional cargo. The reporter recipient HEK293T cells were treated with EVs containing CD81-FKBP, FRB-tTA, and fusogenic protein. As a fusogenic protein, wildtype or mutant (P127D) VSV-G was used. Firefly luciferase (Fluc) was used as a negative control. Donor cells were cultured with 10 nM of rapamycin. N = 3, means ± SD. Data were analyzed by one-way ANOVA followed by Dunnett’s test against non-treatment control. (B) Intracellular tTA release from EVs. Recipient reporter cells were transfected to express TEV protease (TEVp) or mock. Recipient cells were treated with EVs containing either CD81-(TEV)-tTA/VSV-G or CD81-FKBP/FRB-tTA/VSV-G. Donor cells were cultured with 10 nM of rapamycin. N = 3, means ± SD. Data were analyzed by Student’s *t*-test.

EVs fuse with the endosomal membrane by the fusogenic activity of VSV-G and release FRB-tTA into the cytoplasm. However, it is possible that the residual rapamycin tethered FRB-tTA to CD81-FKBP and hampered cytoplasmic release. To investigate the extent of POI release from the EVs, we intentionally cleaved the FKBP domain from CD81 to release FRB-tTA and compared it to the non-cleaved condition. We transfected the recipient cells to express TEV protease (TEVp), which releases the FKBP portion from CD81 by the enzymatic cleavage of the TEVp recognition sequence flanked by CD81 and FKBP. As a control, we used EVs containing CD81-(TEV)-tTA, which can release the tTA domain upon enzymatic cleavage by TEVp and induce the expression of the reporter gene under the TRE promoter^11^. As shown in Fig. 4B, as expected, EVs containing CD81-(TEV)-tTA and VSV-G induced reporter gene expression only in recipient cells expressing TEVp. Without TEVp expression, tTA fused to CD81 and was not released into the cytoplasm. In contrast, EVs containing CD81-FKBP, FRB-tTA, and VSV-G induced reporter gene expression regardless of TEVp expression. This result suggested that FRB-tTA was spontaneously released from CD81-FBKP without enzymatic cleavage and delivered intracellularly.

### Alternative EV-enriched proteins as a tether for FRB-POI

In this study, we chose CD81 as an FKBP-fusion protein to tether FRB-POI for controlled loading into EVs. We switched the CD81 with CD63 or HIV Gag proteins to confirm the general applicability of our strategy. CD63 is an EV marker belonging to the tetraspanin superfamily^20^, whereas HIV Gag is known to be enriched in EVs as cargo protein^21^. We hypothesized that both proteins could serve as FKBP-fusion proteins and recruit FRB-POI in the presence of rapamycin. In the next experiment, FKBP-fused CD63 or Gag were expressed in the donor HEK293T together with FRB-tTA and VSV-G, and concentrated EVs were applied to the recipient cells (Fig. 5). We observed that CD63-FKBP and Gag-FKBP enriched FRB-tTA in the presence of rapamycin and delivered FRB-tTA to the recipient cells, despite weaker induction of reporter gene expression compared to that of CD81-FKBP. Among the EV-enriched proteins tested, CD81 was found to be the best for rapamycin-mediated protein loading and functional delivery to recipient cells.

**Fig. 5.**
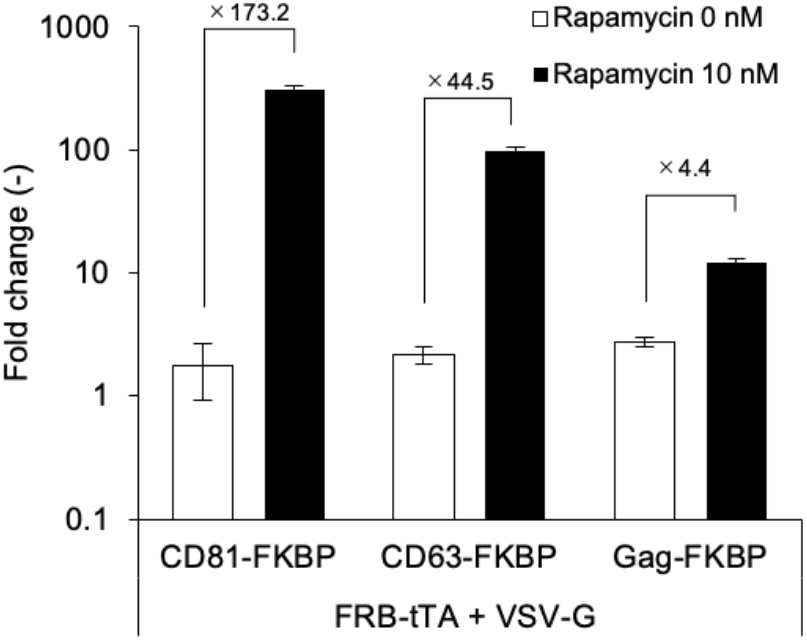
Alternative FKBP-fusion proteins for the delivery of FRB-tTA. Donor HEK293T cells were transfected with plasmids encoding FRB-tTA, VSV-G, and either CD81-FKBP, CD63-FKBP, or Gag-FKBP, and cultured with 0 or 10 nM rapamycin. The concentrated EVs were applied to the recipient cells, and reporter gene expression was measured. Numbers above the bars indicate the fold-increase in reporter gene expression compared to the no rapamycin control. N = 3, means ± SD.

### Delivery of FRB-Cre-HiBiT into recipient cells

Lastly, we switched the POI from tTA to Cre recombinase to verify the versatility of the system. The recipient cells transfected with the reporter plasmid were treated with EVs containing CD81-FKBP, FRB-Cre-HiBiT, and VSV-G. In this assay, the recipient cells initially expressed the red fluorescence protein mKate as a reporter gene and following the cytoplasmic delivery of FRB-Cre-HiBiT, recombination of the LoxP sequence in the reporter plasmid led to the expression of green fluorescent protein EGFP. As shown in Fig. 6, EVs from donor cells treated with 10 nM rapamycin strongly induced EGFP expression in the recipient cells compared to EVs without rapamycin treatment and control, indicating that FRB-Cre-HiBiT was delivered to the recipient cells by the EVs.

**Fig. 6.**
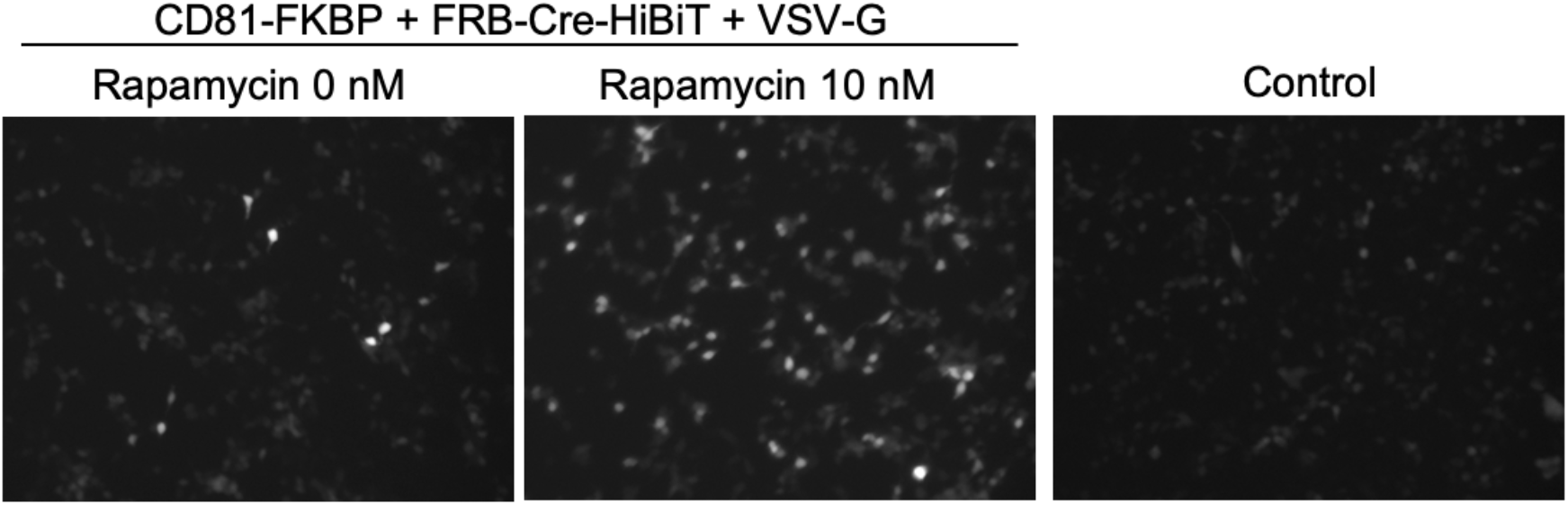
Delivery of FRB-Cre proteins by EVs.

Recipient HEK293T cells were treated with EVs containing CD81-FKBP, FRB-Cre-HiBiT, and VSV-G. After 24 h, the cells were observed under a fluorescence microscope, using the GFP channel.

## Discussion

In this study, we characterized and evaluated rapamycin-induced protein loading into EVs and functional delivery into recipient cells. Efficient and specific loading of EVs remains a challenge for the drug delivery applications of EVs. Multiple loading methods have been described, including electroporation^22^ and physical methods (incubation, permeabilization, freeze-thaw, sonication, and extrusion)^23^. Compared to the conventional methods described in the literature above, loading of POIs inside EV-producing cells is a straightforward approach because EVs are inherently produced by the cells. However, the detailed mechanism of how cells incorporate specific proteins into EVs remains largely unknown, and this leads to difficulties in the efficient loading of POIs in donor cells.

One solution is to utilize protein-protein interactions to recruit POIs into EVs. Sterzenbach et al. reported that the protein-protein interaction between WW tag and Nedd4 family interacting protein 1 (Ndfip1) could be used for loading POI into EVs^24^. Yim et al. achieved efficient loading of POIs into EVs using light-induced protein-protein interaction^25^. The reversible and specific interaction by external stimuli is a feasible way to load POIs into EVs because the POIs can be spontaneously released from EVs. Zhang et al. used split GFP, namely GFP11 and GFP1-10, to achieve the loading of POIs into EVs^18^. They used VSV-G as a fusogenic protein, as in this study, to facilitate membrane fusion in recipient cells and cytoplasmic release of the cargo. Although the POIs fused with GFP1-10 were efficiently encapsulated into EVs, it is unclear how the interaction between GFP1-10-tagged EV protein and GFP11-POI was dissociated and POI was released from EVs inside the recipient cells.

In this study, we utilized the rapamycin-induced interaction of FKBP and FRB, and a similar approach has been reported previously^14,26^. Heath et al. established a loading method for encapsulating FRB-fused Cre into EVs. They demonstrated that EVs themselves could not achieve the intracellular delivery of FRB-Cre; therefore, they identified compounds that could facilitate the endosomal escape of EVs. They revealed that “active loading” (loading FRB-Cre by rapamycin analog) only achieved a slight enrichment of FRB-Cre into EVs (up to 4-fold) compared to “passive loading” (without rapamycin analog). They observed that EVs prepared by both passive and active loading of FRB-Cre achieved an equal extent of functional delivery in the presence of endosomal escape reagents. In contrast, we demonstrated a 70-fold (Fig. 2D) increase in FRB-Cre loading using rapamycin. Although EVs enriched with POIs by active loading can functionally deliver the cargo into the cytoplasm of recipient cells, passive loading of FRB-tTA barely achieved the functional delivery of cargo even in the presence of an endosomal escape mechanism, in this case, VSV-G (Fig. 4A). This discrepancy may be explained by the differences in the quantification methods for POIs inside EVs (highly sensitive HiBiT quantification vs. western blotting), dimerizers (up to 100 nM of rapamycin vs. 500 nM rapalog), assay readouts (luciferase assay vs. fluorescence imaging), culture conditions for donor cells (attachment vs. suspension), and preparation method of EVs (PEG precipitation vs. ultracentrifugation). It should be noted that the EV preparation in the study by Heath et al. contained calnexin, an ER marker that is usually considered as a negative marker for EVs, suggesting that their EV preparations contained contaminants from cellular debris. Compared to their approach using endosome escape-enhancing compounds such as chloroquine or AP21967, we demonstrated that co-expression of the viral fusion protein, VSV-G, significantly increased cargo protein delivery. Our approach appears to be more straightforward because no treatment with toxic endosomal escape reagents is necessary to achieve intracellular delivery of POIs. As shown in Fig. 4B, the cargo proteins fused with FRB domain were spontaneously released from CD81-FKBP inside the recipient cells. This suggested that the preparation process, namely PEG precipitation, diluted rapamycin, and this reduction of rapamycin leads to the spontaneous dissociation of the interaction between FKBP and FRB.

We successfully delivered two different cargo POIs, FRB-tTA and FRB-Cre, to recipient cells. This delivery system depends on the fusogenic activity of VSV-G (Fig. 4A). Conversely, EVs without fusogenic proteins did not achieve successful cytoplasmic delivery of POIs in this study. This result strongly supports the previous findings that EVs generally have a scarce level of fusogenic activity and are thus unable to deliver the cargo into the recipient cells without the exogenous expression of fusogenic proteins^10,11,18,27–29^. Although VSV-G is a potent fusogenic protein that markedly improves the delivery capacity of EVs, it is an immunogenic protein because of its viral origin. Therefore, VSV-G-modified EVs would not be suitable for *in vivo* use, and the use of alternative fusogenic proteins or membrane fusion machinery would be desirable. One study demonstrated that EVs containing mRNA could be modified with a mouse-derived fusion protein called syncytin-A and successfully deliver the cargo to recipient cells^29^. The engineering of EVs with non-VSV-G fusion proteins is a feasible approach for future clinical applications.

Taken together, we successfully developed an efficient loading method controlled by rapamycin that can encapsulate POIs into EVs. The EV-based protein delivery platform holds immense potential for therapeutic delivery of biomolecules. Further engineering should improve the functionality of EVs and expand their delivery capacity.

## Acknowledgments

We extend our gratitude to Yumi Yukawa for providing technical assistance in plasmid construction. We appreciate the helpful discussions with Professor Etsuo A. Susaki and Professor Yumi Kumagai at Juntendo University on the Cre-LoxP reporter system. All illustrations in this study were created using BioRender.com.

This work was supported in part by JSPS KAKENHI (Grant-in-Aid for Early-Career Scientists 18K18386 and 20K15790 to MS), a research grant from the JGC-S Scholarship Foundation (to MS), and the “Dynamic Alliance for Open Innovation Bridging Human, Environment and Materials” (MEXT).

