## Supplementary Tables for "Engineering of extracellular vesicles for small molecule-regulated cargo loading and cytoplasmic delivery of bioactive proteins"

### Supplementary Table 1

| Number | Category | Name | Manufacturer | Cat# |
|---|---|---|---|---|
| 1 | Reagent | Rapamycin | Wako | 184-02531 |
| 2 | NanoLuc substrate | Nano-Glo Luciferase Assay System | Promega | N1120 |
| 3 | NanoLuc substrate | Nano-Glo HiBiT Lytic Detection System | Promega | N3030 |
| 4 | Buffer | RIPA buffer | Nacalai Tesque | 16488-34 |
| 5 | WB detection | ImmunoStar LD | Wako | 296-69901 |
| 6 | Magnet beads | Protein G Mag Sepharose | Cytiva | 28951379 |
| 7 | Antibody | Anti-HiBiT antibody, mouse monoclonal (30E5) | Promega | CS2006A01 |
| 8 | Antibody | Anti-Myc antibody, mouse monoclonal (My3) | MBL | M192-3S |
| 9 | Antibody | Anti-CD81 antibody, mouse monoclonal (17B1) | Wako | 015-27771 |
| 10 | Antibody | Anti-VSV-Glycoprotein, mouse monoclonal (8G5F11) | Merck | MABF2337-25UG |
| 11 | Antibody | Anti-calnexin, mouse monoclonal (4F10) | MBL | M178-3MS |
| 12 | Antibody | TetR Monoclonal Antibody (Clone 9G9) | Clontech | Z1131N |
| 13 | Antibody | anti-GAPDH, monoclonal, HRP-conjugated | Wako | 015-25473 |
| 14 | Antibody | Goat anti-mouse IgG-HRP | TCI | G0407-VIAL |
| 15 | Antibody | Normal Mouse IgG, Whole Molecule, Purified | Wako | 140-09511 |

#### Supplementary Table 2

| Number | Name | Insert | Availability | Reference |
|---|---|---|---|---|
| 1 | pTRE3G-NlucP | C-terminally PEST-tagged NanoLuc under TRE3G promoter | Addgene #162595 | 1 |
| 2 | pcDNA3.1-CD81-(TEV)-tTA | human CD81-TEV cleavage site-tTA | Addgene #162598 | 1 |
| 3 | pcDNA3.1-CD81-(TEV)-Cre | human CD81-TEV cleavage site-Cre recombinase | Addgene #167939 | 1 |
| 4 | pcDNA3.1-TEVp | TEV protease | Addgene #162599 | 1 |
| 5 | pcDNA3.1-LoxP-mKate-LoxP-EGFP | LoxP-flanked mKate and EGFP under CMV promoter | Addgene #167939 | 1 |
| 6 | pCMV-VSV-G-Myc | VSV-G, C-terminal Myc tag | Addgene #80054 | 2 |
| 7 | pCMV-VSV-G(P127D)-Myc | VSV-G(P127D mutant), C-terminal Myc tag | Addgene #80055 | 2 |
| 8 | pCMV-Luc | Firefly luciferase | Addgene #45968 | - |
| 9 | pcDNA3.1-CD81-(TEV)-FKBP-Myc | human CD81-TEV cleavage site-FKBP, C-terminal Myc tag | deposited to Addgene* | This study |
| 10 | pCAG-Gag-FKBP-Myc | HIV Gag-FKBP, C-terminal Myc tag | deposited to Addgene* | This study |
| 11 | pCMV-CD63-FKBP-Myc | human CD63-FKBP, C-terminal Myc tag | deposited to Addgene* | This study |
| 12 | pCMV-FRB-tTA-Myc | FRB-tTA, C-terminal Myc tag | deposited to Addgene* | This study |
| 13 | pcDNA3.1-tTA | tTA | - | This study |
| 14 | pcDNA3.1-FRB-Cre-HiBiT | FRB-Cre, C-terminal-HiBiT tag | deposited to Addgene* | This study |
| 15 | pcDNA3.1-Cre-HiBiT | Cre, C-terminal-HiBiT tag | - | This study |

1 M. Somiya and S. Kuroda, Anal. Chem., vol. 93, no. 13, pp. 5612–5620, Apr. 2021, doi: 10.1021/acs.analchem.1c00339.
2 J. Votteler et al., Nature, vol. 540, no. 7632, pp. 292–295, Nov. 2016, doi: 10.1038/nature20607.
* Will be available through Addgene

DNA sequence of plasmids is available upon request
